# Predicting microscale beat patterns from nanoscale chemomechanics in eukaryotic flagella

**DOI:** 10.1101/2024.08.14.607876

**Authors:** James F. Cass, Hermes Bloomfield-Gadêlha

**Affiliations:** School of Engineering Mathematics and Technology, University of Bristol, Bristol, UK; School of Engineering Mathematics and Technology, and Bristol Robotics Laboratory, University of Bristol, Bristol, UK

## Abstract

We present quantitative predictions for experimental observables—amplitude, frequency and wavelength—of the eukaryotic flagellar beat in terms of underlying molecular chemomechanical parameters. Flagellar beating, an incompletely understood self-organized process arising from the collective action of dynein molecular motors, is modelled as a reaction-diffusion (RD) system with an oscillatory instability arising from motor-induced microtubule sliding. While the RD model accurately reproduces beating patterns of bull spermatozoa and *C. Reinhardtii*, existing linear analyses and simulations are unable to provide a complete framework for understanding nonlinear waveform formation. Here, we derive analytical expressions that reveal the nonlinear dependence of beat characteristics on parameters such as motor binding duty ratio, stepping velocity, and axonemal resistance. Our analysis uncovers a novel out-of-equilibrium mechanism for base-to-tip wave propagation, involving an interference pattern between unstable standing wave modes that generates travelling waves. Predicted beat patterns agree remarkably with numerical simulations, even far from the critical point marking the onset of oscillations. This unveils key molecular parameters that govern oscillation initiation, amplitude saturation, frequency shifts, and the spatial phase gradient crucial for generating propulsive hydrodynamic force. Our results yield biophysical understanding of how molecular interactions shape flagellar beating patterns, allowing for the inference of molecular properties from macroscopic observations. This challenges existing hypotheses on wave generation and demonstrates the power of nonlinear analysis to uncover new phenomena beyond the reach of linear models and computational studies alone.

## I. INTRODUCTION

The beating patterns of eukaryotic flagella and cilia, with typical lengths ∼10 – 50 µm, arise from selforganized processes within the cyclindrical structure of doublet microtubules known as the *axoneme* [1, 2] (see Fig. 1**a**). Dynein motor proteins, roughly three orders of magnitude smaller at the scale of ∼10 nm, generate active sliding forces between doublets that, remarkably, are coordinated in space and time to bend the axoneme in periodic waves [3, 4]. This oscillatory behavior plays a crucial role in locomotion [5, 6] and transport processes [7] across various microbiological systems. Experimental studies have revealed a multitude of flagellar waveforms displaying a range of amplitudes, frequencies and wavelengths [8], with both base-to-tip and tip-to-base wave propagation [9], and with hydrodynamic synchronization entraining metachronal waves that propagate over many cilia [10, 11]. However, a theoretical framework that directly predicts the emergent waveform (as characterised by its amplitude, frequency and phase distribution along the flagellum) from underlying chemomechanical mechanisms is still lacking.

**FIG. 1.**
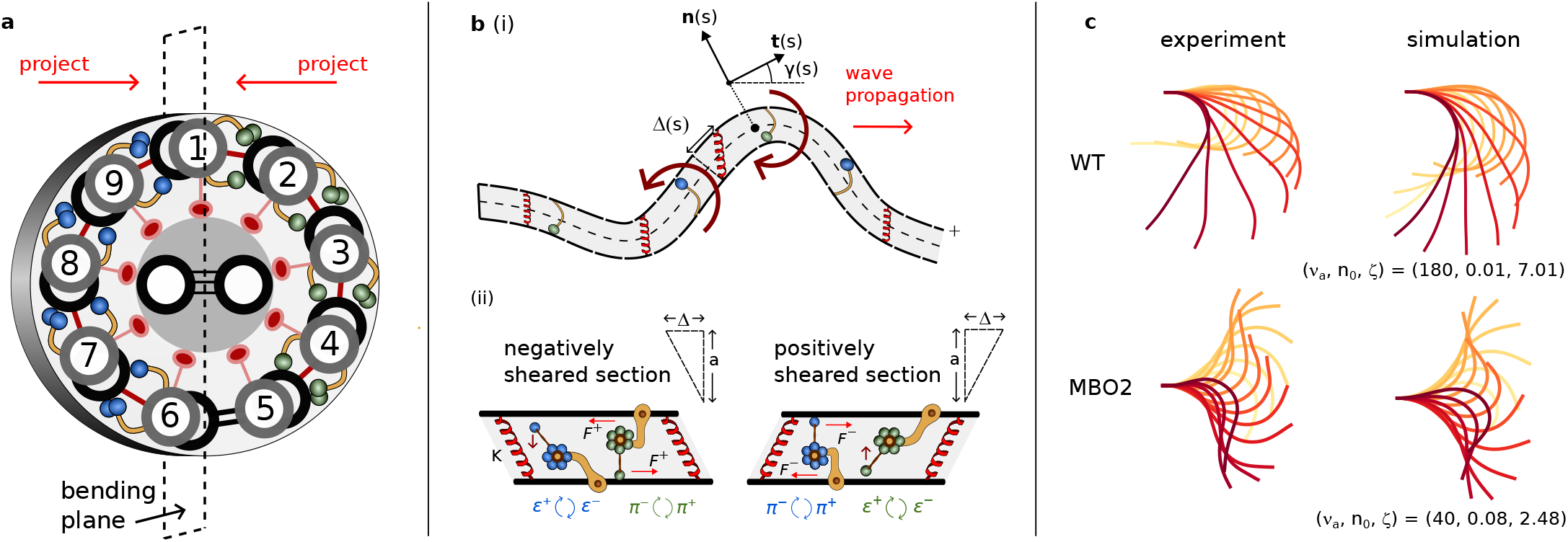
Reaction-diffusion modelling framework (figure adapted from [12]). **a** Schematic of a cross section of the axoneme, consisting of 9 doublet microtubules arranged cylindrically and connected by active dynein molecular motors and passive elastic cross linkers. **b** (i) The flagellum is modelled as a two-filament viscoelastic rod with shear and bending resistance, with configuration characterised by the local shear angle *γ*(*s, t*) (ii) sections of positive and negative sliding displacements Δ driven by dynein motors engaged in a local tug-of-war exerting forces *F*^*±*^, with attachment rates *π*^*±*^ and detachment rates *ϵ*^*±*^. **c** Simulations of the reaction-diffusion model using fitted parameters [12] to experimental data of wild-type (WT) (*R*^2^ = 0.88 *±* 0.044) and MBO2-mutant *C. Reinhardtii* (*R*^2^ = 0.967 *±* 0.012) ciliary beating patterns (using data from [13]). Key theoretical results concerning the nonlinear dynamics of these large amplitude oscillations are the focus of this study.

Biophysical modelling has mainly focused on the idea that mechanical stresses on dynein due to flagellar deformations may regulate motor activity [14–16]. Modelling studies have shown self-sustained oscillations can be generated whether the mechanical feedback is due to forces tangential or normal to the axoneme or due to imposed curvature [15, 17–20]. An alternative, open-loop, mechanism suggests that dynein regulation is not required, and that oscillations are instead driven by a flutter instability arising from the axial forces imparted by steady dynein activity [21–24]. Studies have analyzed flagellar dynamics using elastohydrodynamic approximations and numerical simulations [14, 15, 20, 25–27], examined the linear onset of oscillations near the Hopf bifurcation [17, 20, 28], revealed generic properties of self-organised flagellar oscillations [17, 29] or proposed phenomenological phase oscillator models [30, 31]. While providing fundamental physical insights, most approaches lack an explicit connection between experimental observables (typically measurements of the flagellar centerline, e.g. [28, 32]) and nanoscale chemomechanical properties of the flagellar-dynein interaction (aside from a limited range of parameters explored in simulations) due to the complexities involved in finding solutions to high-order partial differential equation models beyond the regime of linear stability. As a result, open questions remain regarding how molecular motor properties modulate flagellar waveforms and swimming gaits within these model systems that involve many coupled equations. For example, how does the duty ratio of molecular motor binding or the velocity of motor stepping affect the mechanical output of the flagellum?

Here we address this gap through a combined analytical and computational study of a model flagellar system that accurately reproduces experimental waveforms (Fig. 1**c**). In previous work [12], we considered the low viscosity, large-amplitude regime of sliding-controlled (i.e. dynein regulation by forces *tangential* to the doublet microtubules) flagellar beating, where the balance of forces and torques on the swimmer is decoupled from the internal moment balance responsible for generating bending deformations. In this regime, external hydrodynamic resistance has a negligible effect on the beating patterns, consistent with experimental evidence in distinct eukaryotic species [33–35]. Consequently, an intrinsic model for beat patterning can be derived, culminating in a system of reaction-diffusion equations that couple shearing and bending of the passive elastic axoneme with active molecular motor forces and dissipation. The view this perspective offers is of chemomechanical pattern formation that emerges from the interactions of linearly coupled shear oscillators distributed along the flagellum [36, 37] (see Fig. 1**b**(ii)), connecting flagellar beating with well studied oscillatory pattern forming systems such as the Belousov-Zabotinsky chemical reaction [38, 39], which occur in a fluid rather than solid medium. In this picture, synchronous oscillations along the flagellum correspond to time-reversible standing waves that do not generate a propulsive force at low Reynolds number but, importantly, relative phase differences between neighbouring oscillators can be generated that engender net hydrodynamic force that can be harnessed for swimming or fluid pumping.

A significant finding of our previous work was that there exist parameters of the reaction-diffusion model that accurately fit the beating patterns of *C. Reinhardtii* (see Fig. 1**c**)—suggesting a modification of the conclusion (from linear analyses) that sliding-controlled oscillations can not produce the base-to-tip waves characteristic of short-ciliated eukaryotes [13, 35]. In order to understand the phenomenon of base-to-tip propagation in sliding-controlled models it is therefore necessary to explore the nonlinear dynamics of large amplitude flagellar beating, going beyond foundational linear analyses [17, 28]. In this paper, by performing a weakly nonlinear expansion around the oscillatory instability of the reaction-diffusion model of flagellar beating we derive analytical expressions for waveform characteristics in terms of motor binding kinetics, protein friction and elasticity. The analysis provides testable predictions for how waveform shape, frequency, propagation direction and propulsive force generation depend on dynein activity, coordination, and characteristic length and time scales beyond the regime of linear stability. A key result is the *emergence* of baseto-tip wave propagation through the novel interaction of multiple unstable standing wave modes—time-reciprocal standing wave solutions near equilibrium are modulated by shorter wavelength modes with a relative temporal phase difference as dynein activity is increased.

The following section provides a brief introduction to the reaction-diffusion model of flagellar beating [12]. We proceed to our results, discussing waveform modulations as the system is driven further from equilibrium characterised by the saturating amplitude, a shifting frequency and the development of a spatial phase gradient. Finally we explore far from equilibrium oscillations using numer-ical simulations. The analytical and numerical methods by which the results were obtained are detailed in Appendix A and Appendix C and code is provided at github.com/polymaths-lab. We conclude with a summary and discussion of the biophysical implications of these results.

## II. REACTION-DIFFUSION MODEL

The flagellar reaction-diffusion model is derived in detail in previous work [12]. Briefly, the governing equations for a two-filament flagellum of length *L* and diameter *a* read,

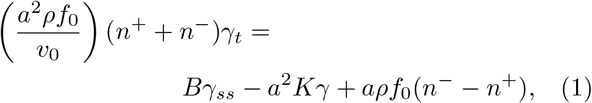

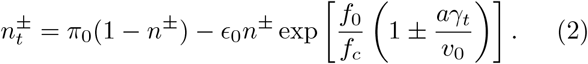

where *γ*(*s, t*) is the flagellum shear angle, or equivalently the tangent angle relative to the orientation at *s* = 0 (see Fig. 1**b** (i)) and *n*^*±*^(*s, t*) are the proportions of bound motors in an element of size *ds* on the plus or minus filaments (subscripts *s* and *t* denote space and time derivatives). Eq. (1) represents the internal balance of moments in the low-viscosity limit[40] with the bending rigidity *B* and shear stiffness per unit length *K* characterising the elastic resistance of the flagellum to bending and shearing respectively.

The motor force is modelled by *F*^*±*^ = *f*_0_(1 ± Δ_*t*_*/v*_0_) where *f*_0_ is the force of a stalled motor and *v*_0_ is the unforced velocity of a motor. Here, the sliding displacement Δ(*s, t*) = *aγ*(*s, t*) (Fig. 1**b**) is the difference in arclength between the plus and minus filaments at a cross section. Molecular motors lie along the filaments with linear density *ρ*, exerting antagonistic active moments *aρn*^*±*^*F*^*±*^ in a tug-of-war between the filaments. The two-state motor kinetics are expressed by Eq. (2), where the constants *π*_0_ and *ϵ*_0_ are the base attachment and detachment rates of motors. The detachment rate is modulated exponentially by the ratio of sliding forces *F*^*±*^ felt by the motors to a characteristic unbinding force *f*_*c*_, implementing sliding-controlled feedback in the motor dynamics.

The boundary conditions we consider enforce that sliding of the filaments is constrained at the base, *γ*(0) = 0, and that there are no external moments at the distal end, *Bγ*_*s*_(*L*) = 0. Eq. (1) and Eq. (2) with the boundary conditions form a closed system of three partial differential equations that describe self-sustained beat pattern generation in low viscosity, decoupled from the equations of motion of the swimmer in the fluid medium.

We cast the system in terms of non-dimensional parameters using the length scale *L* of the flagellum, and time scale *τ* ^*b*^ = (*π*_0_ + *ϵ*_0_ exp(*f*_0_*/f*_*c*_))^−1^ of motor binding relaxation.

***Activity:*:** *ν*_*a*_ = *ρf*_0_*/*(*aK*). The characteristic motor force ∼ *aρf*_0_ relative to passive shear resistance ∼ *a*^2^*K*. The parameter *µ*_*a*_ = *aρf*_0_*L*^2^*/B* has been more commonly used as a non-dimensional activity parameter in previous studies [20, 25, 26]. The parameter *ν*_*a*_ will be the primary bifurcation/control parameter in our study.

***Diffusivity:*:** 𝒟 = *B/*(*a*^2^*KL*^2^). The bending resistance ∼ *B/L*^2^ relative to the shear stiffness ∼ *a*^2^*K*. The inverse parameter *µ* = 𝒟^−1^ has been commonly used in previous studies. For *C. Reinhardtii* we can estimate 𝒟 ≈ (840 pN µm^2^)*/*(80 pN rad^−1^)(10 µm)^2^ ≈ 0.1 using experimental measurements in [41].

***Duty ratio:*:** *n*_0_ = *π*_0_*τ* ^*b*^. The duty ratio of the system when in equilibrium (locked in the tug-of-war) i.e. the fraction of time spent bound by the motors.

***Internal friction:*:** *ζ* = *a/*(*v*_0_*τ* ^*b*^). The diameter of the axoneme *a*, relative to the characteristic step length of an unforced motor *v*_0_*τ* ^*b*^. This can be seen as a non-dimensionalised internal drag coefficient of the dynein motors due to the dependence on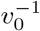.

*f*^*^ = *f*_0_*/f*_*c*_: The ratio of the motor stall force to the characteristic unbinding force. In this paper we only consider *f*^*^ = 2 to reduce complexity [28].

E = *ξ*^*n*^*L*^2^*/a*^2^*Kτ* ^*b*^: The ratio of hydrodynamic forces ∼ *ξ*^*n*^*L*^2^*/τ* ^*b*^ (where *ξ*^*n*^ is a hydrodynamic drag coefficient per unit length) to elastic shear forces ∼ *a*^2^*K*. Here we consider ℰ ≪ 1, neglecting any effect of hydrodynamic resistance on the flagellar beat pattern formation. For example, for *C. Reinhardtii* in water we can estimate ℰ ≈ (10^−3^ pN s µm^−2^)(10 µm)^2^(50 Hz)*/*(80 pN rad^−1^) ≈ 0.06 [13, 41], where we have used the approximate beating period in place of the binding time *τ* ^*b*^.

In the above we have given convenient names to the four key parameters (*ν*_*a*_, 𝒟, *n*_0_, *ζ*) that will be the studied in the following. Typical parameter values from the literature and corresponding non-dimensional parameter ranges are given in Tab. I. In terms of these parameters, Eq. (1)-(2) become the reaction-diffusion type system,

**TABLE 1.** Typical values of dimensional and nondimensional parameters. Adapted from the parameter values given in [20]. Since the motor binding time *τ*^*b*^ is unknown, we use typical ciliary beating periods;Fig. 3**b** suggests the beating frequency is set by the inverse binding time.

| dimensional | nondimensional |  |
| --- | --- | --- |
| $L$ 10–50 $\mu\text{m}$ | flagellar length scale ($L$) | |
| $a$ 0.2 $\mu\text{m}$ | sliding length scale ($a$) | |
| $K$ $2 \times 10^3$ pN $\mu\text{m}^{-2}$ | force scale ($a^2 K$) | |
| $\tau^b$ 20–100 ms | binding time scale ($\tau^b$) | |
| $B$ 90–1700 pN $\mu\text{m}^2$ | $\mathcal{D} = B/a^2 L^2 K$ | 0.01–0.2 |
| $\xi^n$ $10^{-3}$–1 pN s $\mu\text{m}^{-2}$ | $\mathcal{E} = \xi^n L^2/a^2 K \tau^b$ | 0.01–1500 |
| $f_0$ 1–5 pN | $\nu_a = \rho f_0/aK$ | 2.5–12.5 |
| $f_c$ 0.25–2.5 pN | $f^* = f_0/f_c$ | 2 |
| $v_0$ 5–7 $\mu\text{m s}^{-1}$ | $\zeta = a/v_0 \tau^b$ | 0.29–2 |
| $\rho$ $10^3$ $\mu\text{m}^{-1}$ | | |

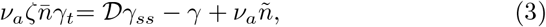

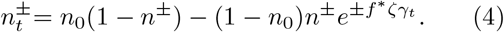

where we have introduced the total number density of attached motors 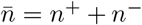 and the differential motor binding density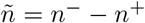.

We will continue to use the symbols *s* for the arclength and *t* for time to describe our results, but the reader should take these to imply the non-dimensional versions *s/L* and *t/τ* henceforth.

## III. RESULTS

The combination of weakly nonlinear analysis (see Appendix A) and numerical simulations (Appendix C) provide insights into the modulation of flagellar waveforms as the reaction-diffusion system (Eq. (3) and Eq. (4)) moves away from its point of instability, known to be an oscillatory Hopf bifurcation from previous work [12]. To apply perturbation theory, the relevant small parameter is the relative distance from the bifurcation point, defined by setting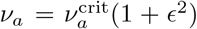, where 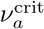 marks the level of motor activity required at the onset of oscillations. The symmetry of the problem, taking *γ* → −*γ* and *n*^−^ ↔ *n*^+^, causes various terms to vanish in the perturbation expansion, simplifying the analysis (Appendix A).

In the following, we present analytical expressions describing modulations of the beat as the system is driven away from equilibrium (i.e. in powers of *ϵ*). The key results appear at 𝒪 (*ϵ*^3^), where a spatially-dependent phase *ϕ*(*s*) emerges that modulates the standing waves that appear at orders 𝒪 (*ϵ*) and 𝒪 (*ϵ*^2^). Reciprocal standing waves, like Purcell’s scallop [5], do not lead to a propulsive force that would engender progressive motion, so the emergence of a phase gradient is critical for swimming [17]. The weakly nonlinear solutions obtained then enable us to calculate the hydrodynamic force generated by the emergent non-reciprocal beating patterns.

### A. Oscillatory Instability

Oscillations occur for positive values of the critical expression,

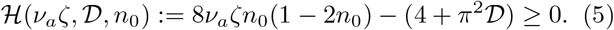

The critical point, ℋ = 0, marks the Hopf bifurcation, or the point of instability of the straight equilibrium configuration *γ* = 0. Eq. (5) should be understood as the difference between an active term, which grows linearly with activity *ν*_*a*_ and internal friction *ζ* and has non-monotonic dependence on the duty ratio *n*_0_, and a passive term including shear resistance (the origin of the numerical 4) and bending resistance (from the term *π*^2^𝒟). The combination *ν*_*a*_*ζ*, that also appears on the left hand side of Eq. (3), can be interpreted as the ratio of the viscoelastic relaxation timescale of the flagellum in tugof-war *τ* ^*f*^ = *ρf*_0_*/Kv*_0_ to the motor binding relaxation timescale *τ* ^*b*^ = (*π*_0_ + *ϵ*_0_ exp(*f*^*^))^−1^.

The critical frequency at the onset of oscillations is given by 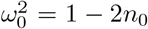 [12]; hence a necessary condition for oscillations is that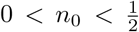. Fig. 2 shows how the unstable interval of *n*_0_ values widens for increasing timescale separation *ν*_*a*_*ζ*. At the midpoint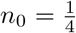, where ℋ is maximal, the critical expression takes the simpler form ℋ^max^(*ν*_*a*_*ζ*, 𝒟) = *ν*_*a*_*ζ* − (4 + *π*^2^𝒟), so that for small values of diffusivity 𝒟 we have the rule of thumb that the timescale separation should be at least a factor of 4 for oscillations to occur. Note that the critical frequency *ω*_0_ is independent of *ν*_*a*_*ζ* (and 𝒟), indicating that close to criticality the frequency is determined only by the dynein binding and unbinding cycle, through the duty ratio *n*_0_.

**FIG. 2.**
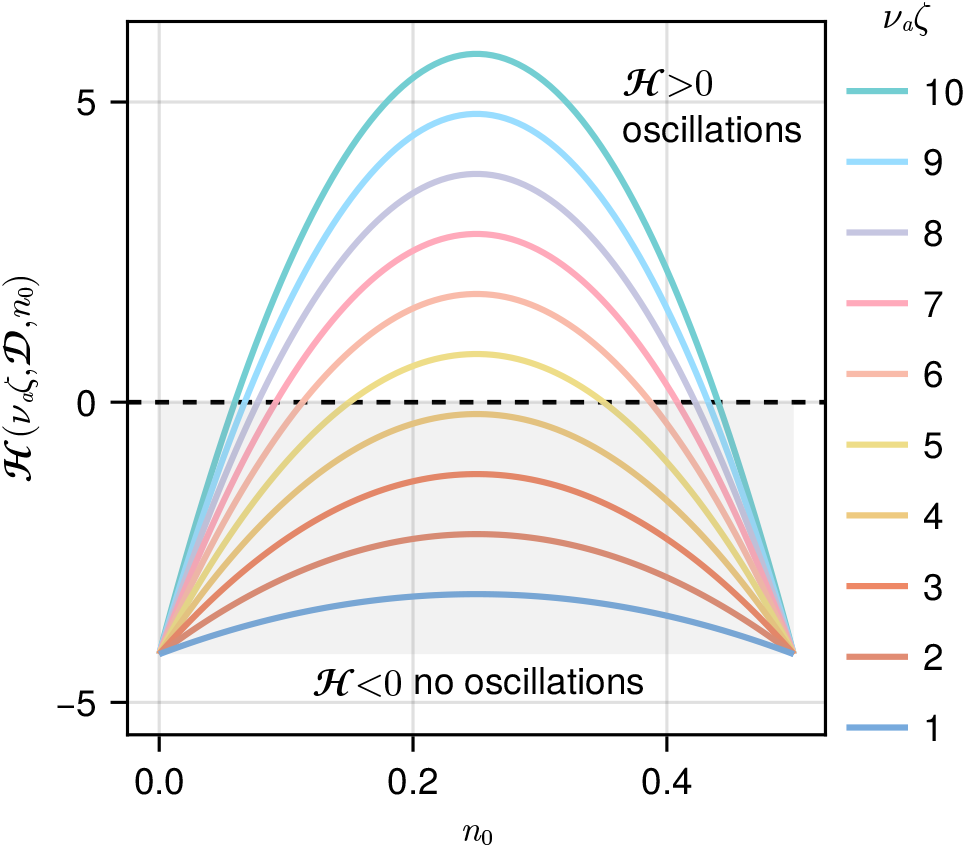
Widening critical region for increasing timescale separation (the ratio of characteristic viscoelastic flagellar relaxation and motor binding times *ν*_*a*_*ζ* = *τ* ^*f*^ */τ* ^*b*^), as a function of the loaded duty ratio *n*_0_ (with 𝒟 = 0.02). The straight equilibrium state *γ*(*s*) = 0 becomes unstable to limit cycle oscillations for values of *n*_0_ such that ℋ *>* 0, where *H*(*ν*_*a*_*ζ*, 𝒟, *n*_0_) is given in Eq. (5).

### B. Standing wave solutions

The solutions to the linear 𝒪 (*ϵ*) equations are *standing waves*, with shear angle dynamics given by

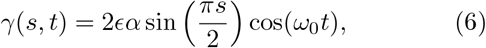

with a saturation amplitude of the shear *γ*^sat^ := 2*ϵα* at *s* = 1 that depends on the parameters according to

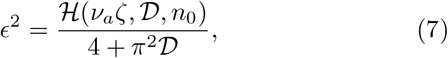

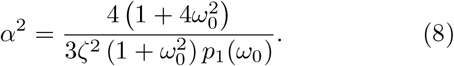

where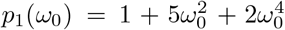. It will often be con-venient to write expressions in terms of *ω*_0_ rather than the duty ratio *n*_0_ to avoid a proliferation of square roots. The expression on the right of Eq. (8) is always positive; thus the Hopf bifurcation is *supercritical* for all parameter values[42]. The denominator introduces dependence on *ζ*, independently of *ν*_*a*_, in contrast to the critical expression.

Before discussing the dependencies implied by Eq. (7) and Eq. (8) of the saturation amplitude *γ*^sat^ on the chemomechanical parameters we introduce a revealing decomposition of the solutions into a basis of standing wave modes that derives from the perturbation expansion and is used for numerical simulations (see Appendix C).

### C. Spectral decomposition into standing wave modes

We decompose the beating pattern, in a manner suggested by the perturbation expansion (Appendix A), into a basis of standing waves consisting of products of spatial and temporal Fourier 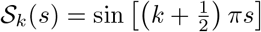 and 𝒯_*l*_(*t*) = exp(*liωt*) for *k, l* = 0, 1, 2, … (various decom-positions are shown below in Fig. 7**c**,**d**). The shear angle *γ*(*s, t*) can be written as a sum of these modes,

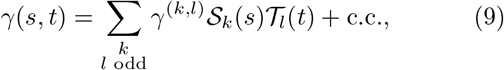

where the coefficients *γ*^(*k,l*)^ are complex numbers in general and c.c. denotes the complex conjugate expression. The double sum in Eq. (9) only runs over odd values of *l* because all even contributions are zero by symmetry. The value of *k* can be understood as the number of interior nodes for *s* ∈ (0, 1) of each spatial mode. The expansion is completely determined by the constants

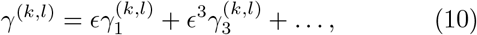

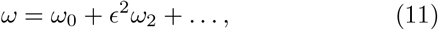

which are expanded in the bifurcation parameter *ϵ*. By truncating the series at various orders in *ϵ* we will examine modulations of the beating pattern due to weak nonlinearity. The linear standing wave solution, Eq. (6), contains only a contribution from the (*k, l*) = (0, 1) mode, where *γ*^(0,1)^ = *ϵα* + 𝒪 (*ϵ*^3^).

In the following, many quantities arise that need to be clearly distinguished notationally. We will use the following conventions:

- Theoretical results use a lower case letter (e.g. *γ*(*s, t*), *ñ*(*s, t*)). Corresponding calculations from simulations use an upper case letter (e.g. Γ(*s, t*), *Ñ*(*s, t*)).
- Space-time standing wave modes are indexed by a tuple (*k, l*), referring to mode 𝒮_*k*_(*s*) 𝒯_*l*_(*t*) for an odd expansion (*γ* or *ñ*), and 𝒞_*k*_(*s*) 𝒯_*l*_(*s*) for an even expansion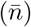. The (complex) coefficient of a mode is indicated by a superscript, for example *γ*^(0,1)^ or Γ^(0,1)^.
- The coefficient functions of *temporal* modes T_*l*_(*t*) are denoted by a single superscript e.g. *γ*^*l*^(*s*) = ∑_*k*_ *γ*^(*k,l*)^ 𝒮_*k*_ (*s*).
- The coefficients of modes can be further decomposed into an asymptotic series in powers of *ϵ*. We denote the coefficient of *ϵ*^*m*^ in the expansion *γ*^(*k,l*)^ by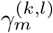. For example 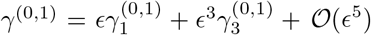.

Hence we use the symbol *γ* for many quantities; the signature: *γ*(*s, t*), *γ*^(*k,l*)^, *γ*^*l*^(*s*), Γ^*l*^(*s*), 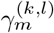 makes clear which is being referred to.

We note here that simulations were carried out using a spectral numerical method that makes use of the expansion into modes, Eq. (9), which reduces taking space or time derivatives to multiplication. Accurate periodic solutions were obtained using a bespoke predictor-corrector scheme described in Appendix C.

### D. Saturation amplitude

The variation of the saturation amplitude *γ*^sat^, a key experimental observable, with the four system parameters is shown in Fig. 3**a**, where each non-varying parameter is fixed at (*ν*_*a*_, 𝒟, *n*_0_, *ζ*) = (5, 0.02, 0.1, 2)—a representative waveform within the physical parameter ranges presented in Tab. I. The lower *x*-axis marks the parameter which is varied while the upper *x*-axis shows the corresponding measure of the distance from the bifurcation *ϵ*.

**FIG. 3.**
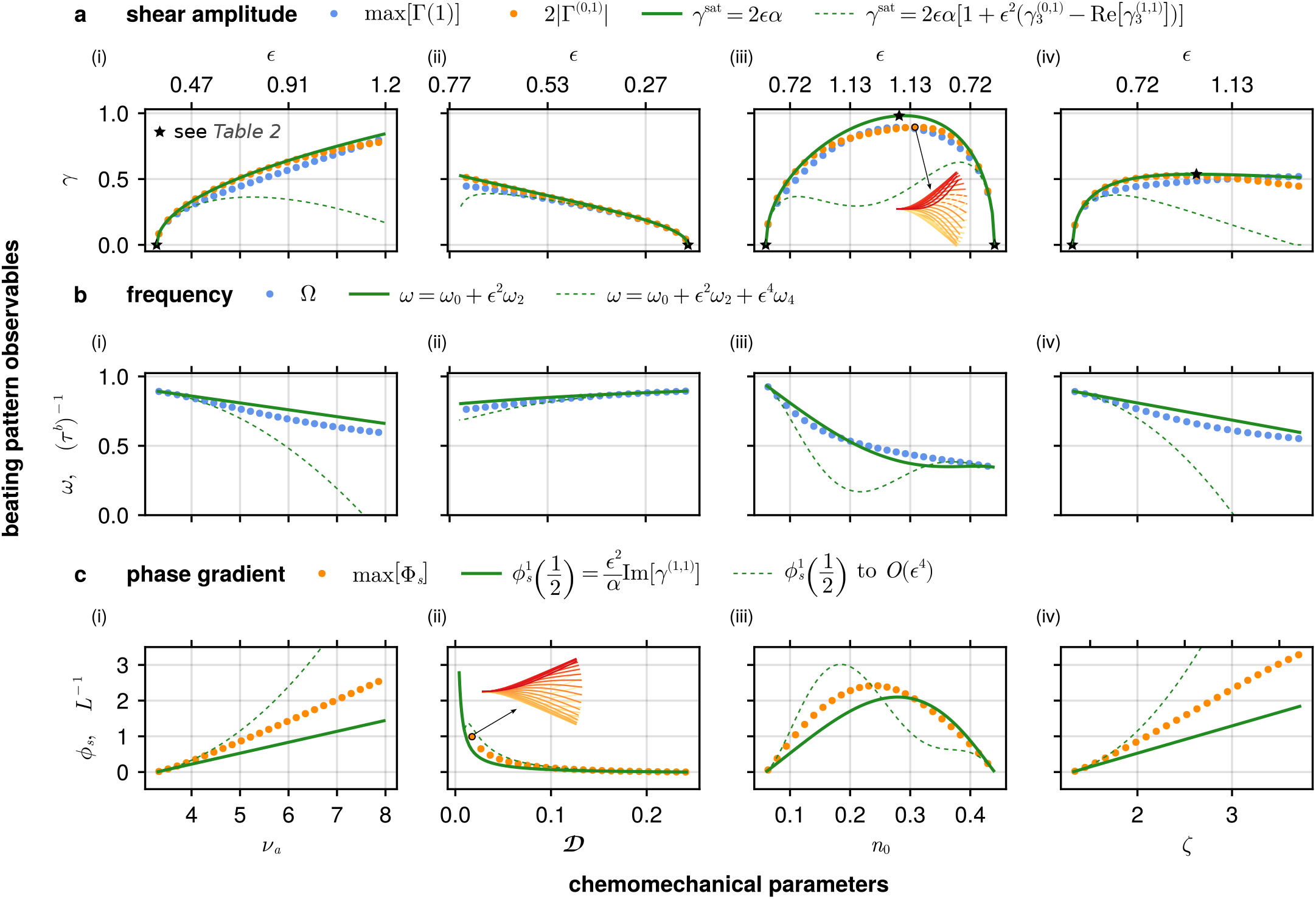
Weakly nonlinear predictions (solid green lines) of key waveform observables of the RD model (amplitude, frequency and phase) as functions of nondimensional chemomechanical parameters (i) activity *ν*_*a*_, (ii) diffusivity 𝒟, (iii) duty ratio *n*_0_ and (iv) internal friction *ζ*, compared with numerical simulations (blue/orange markers). Blue markers are quantities calculated directly from simulations, while orange markers are calculated from the spectral decomposition (Eq. (9)) of the simulation. Dashed green lines show predictions (generally only accurate for small values of *ϵ*) of one order higher in *ϵ* than solid lines. **a** Amplitude of the theoretical saturating shear angle *γ*^sat^ (Eq. (6)) compared with the numerical maximum shear amplitude max[Γ(*s* = 1)] and mode 𝒮_0_(*s*) 𝒯_1_(*t*) amplitude 2|Γ^(0,1)^|. **a**(iii, inset) The beating pattern in the *x*-*y* plane corresponding to the maximum shear amplitude as a function of *n*_0_. Starred points have analytical expressions given in Tab. II. **b** Theoretical frequency *ω* (Eq. (12)), and numerical frequency Ω = 2*π/T* (Appendix C). **c** Theoretical spatial derivative of the phase of oscillations *ϕ*_*s*_ (Eq. (14)), and numerical Φ_*s*_ (from the temporal Fourier transform), evaluated at the midpoint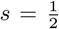. **c**(ii, inset) The beating pattern in the *x*-*y* plane of the largest simulated phase gradient as a function of 𝒟. Parameters are fixed at (*ν*_*a*_, 𝒟, *n*_0_, *ζ*) = (5.0, 0.02, 0.1, 2.0) if not explicitly varying.

We first make some general remarks applicable to all four parameter sweeps in Fig. 3**a**. The solid green lines show the leading order asymptotic prediction *γ*^sat^ = 2*ϵα*, compared with the maximum tip amplitude at *s* = 1 obtained from numerical simulations (denoted max[Γ(1)], blue circles). Remarkably, we find that the prediction is close to the maximal tip shear for a large range of parameter values, even for values of the perturbation parameter *ϵ* greater than unity, a key result of this paper.

The alternative numerical estimate of the saturation amplitude, Γ^sat^ = 2|Γ^(0,1)^|, that comes directly from the spectral decomposition of the simulation, is shown in the same figure (orange circles). Comparing again with the leading order prediction *γ*^sat^ = 2*ϵα*, we see a yet closer agreement than was obtained with the maximal shear max[Γ(1)]; this could be expected since the maximum may combine the effects of many unstable modes when the system is far from equilibrium. We will discuss higher order contributions below, save for a remark here that the *O*(*ϵ*^3^) correction to *γ*^sat^ (dashed green lines) highlights a common feature of asymptotic series [43]. While the approximation converges in the limit *ϵ* → 0, for which the 𝒪 (*ϵ*^3^) approximation indeed becomes closer to the true value, taking more terms in the series at a *fixed* value of *ϵ* does not provide more accuracy [44]. From Fig. 3**a**, we see that the optimal truncation for *ϵ* ≳ 0.2 is the leading term only.

We now discuss the variation of individual parameters in Fig. 3**a**(i)-(iv). The amplitude shows square root growth from the critical point, as is expected for a Hopf bifurcation, growing like (i) 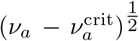 for increas-ing activity *ν*_*a*_ and (ii) 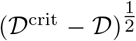 for decreasing dif-fusivity 𝒟. While the amplitude continues to increase with increasing *ν*_*a*_, at 𝒟 = 0 the amplitude reaches a maximum value—an unphysical solution representing the shear oscillations of isolated sections which are uncoupled and unconstrained by the boundary conditions, as discussed previously [12]. The amplitude dependence on the duty ratio *n*_0_ in panel (iii) resembles the quadratic dependence of ℋ on *n*_0_ in Fig. 2, with critical points 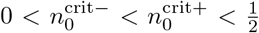. For values close to the critical point, however, the curve is steeper thanFig. 2 (due to the square root), and the peak is skewed to values 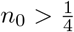 by multiplication with *α*, a monotonically increasing function of[ineq[. The beating pattern in the *x*-*y* plane corresponding to this peak is shown inset in panel (iii). Finally, in panel (iv) we see that the amplitude also displays a non-monotonic dependence on internal friction *ζ*, with a maximum value predicted to be *ζ*^max^ = 2*ζ*^crit^. Given the accuracy of these predictions, we summarise the approximate or exact predicted values for the critical points and extrema (marked with stars in Fig. 3**a**) of these curves in Tab. II, which may be useful for inferring chemomechanical parameters from experimental measurements.

**TABLE 2.** Analytical expressions for the starred points in Fig. 3**a**, relating the shear amplitude *γ*^sat^ = 2*ϵα* to the nondimensional parameters (*ν*_*a*_, 𝒟, *n*_0_, *ζ*). The critical frequency is given by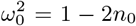. The maximum 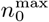 is a linear approximation to the root of ∂*γ*^sat^*/*∂*n*_0_ around *n*_0_ = 1*/*4.

| Point | Expression | Comments |
| --- | --- | --- |
| $\nu_a^{\text{crit}}$ | $\frac{4+\pi^2\mathcal{D}}{4\zeta\omega_0^2(1-\omega_0^2)}$ | Oscillations for $\nu_a > \nu_a^{\text{crit}}$ |
| $\mathcal{D}^{\text{max}}$ | 0 | No bending resistance/only shear resistance |
| $\mathcal{D}^{\text{crit}}$ | $\frac{4}{\pi^2}(\nu_a\zeta\omega_0^2(1-\omega_0^2)-1)$ | Oscillations for $\mathcal{D} < \mathcal{D}^{\text{crit}}$ |
| $n_0^{\text{crit}\pm}$ | $\frac{1}{4}\left(1 \pm \sqrt{1 - \frac{4+\pi^2\mathcal{D}}{\nu_a\zeta}}\right)$ | Oscillations for $n_0^{\text{crit}-} < n_0 < n_0^{\text{crit}+}$ |
| $n_0^{\text{max}}$ | $\approx \frac{1}{4} + \frac{13(\nu_a\zeta-4-\pi^2\mathcal{D})}{16(17\nu_a\zeta-20-5\pi^2\mathcal{D})}$ | $\frac{1}{4} < n_0^{\text{max}} < n_0^{\text{crit}+}$ |
| $\zeta^{\text{crit}}$ | $\frac{4+\pi^2\mathcal{D}}{4\nu_a\omega_0^2(1-\omega_0^2)}$ | Oscillations for $\zeta > \zeta^{\text{crit}}$ |
| $\zeta^{\text{max}}$ | $2\zeta^{\text{crit}}$ | $\gamma^{\text{sat}} \sim \zeta^{-\frac{1}{2}}$ for $\zeta > \zeta^{\text{max}}$ (also see Fig. 8a) |

### E. Frequency shift

The frequency *ω* decreases as the system is moved away from equilibrium in the parameters *ν*_*a*_, 𝒟 and *ζ* (see Fig. 3**b**(i), (ii) and (iv)), as predicted by the negative correction term at second order in *ϵ*.

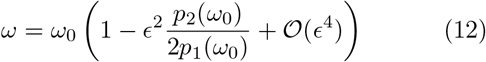

where 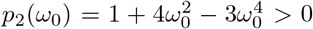 for *ω*_0_ ∈ (0, 1). The dependence of frequency on the duty ratio *n*_0_ is notable, since the monotonic decrease in *ω* for increasing *n*_0_ contrasts with the variation of the amplitude (Fig. 3**a**) which rises and then falls (compare **a**(iii) and **b**(iii)). The prediction *ω* = *ω*_0_ + *ϵ*^2^*ω*_2_ given by Eq. (12) is once again remarkably accurate for values of *ϵ* much larger than one might expect. As with Fig. 3**a** we show the next order correction to the frequency (O(*ϵ*^4^)) which provides a better prediction in the limit *ϵ* → 0, but deviates thereafter.

### F. Phase gradient-driven bending waves

The reciprocal standing waves obtained so far, like Purcell’s scallop [5], do not lead to a propulsive force that would engender progressive motion. By transforming the decomposition, Eq. (9), into the more physical amplitude-phase form

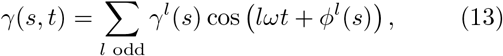

we see that, crucially for swimming, a nonzero phase gradient,

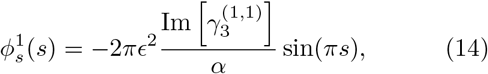

of the fundamental oscillatory mode arises at the next order that enables propulsive force generation[45]. Fig. 3**c** shows the magnitude of the phase gradient at the midpoint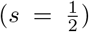, where Eq. (14) is maximal, evaluating to

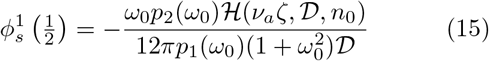

in terms of the four chemomechanical parameters. The wavelength of the beat can be defined via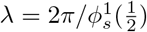. Since *p*_1_, *p*_2_ *>* 0 for *ω* ∈ (0, 1], we see that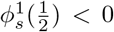, indicating base-to-tip wave propagation for all parameter values. This is in contrast to solutions of linearised sliding-controlled elastohyrodynamics (where hydrodynamic moments are retained), in which the direction of propagation depends on the boundary conditions [17, 19]. The quantity 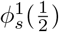 has a simple linear dependence on activity *ν*_*a*_ and internal friction *ζ* (panels (i) and (iv)) through the dependence on the critical value ℋ, Eq. (5), and non-monotonic dependence on *n*_0_ (panel (iii)). Of particular note in Eq. (15) is the dependence on 𝒟, the nondimensionalised bending elasticity or, from the oscillator perspective, the coupling between neighbouring shearable elements. The amplitude of the phase difference scales with 𝒟^−1^, meaning that weak coupling between elements enhances the ability to generate non-reciprocal waveforms. This is realised in the sharp growth in phase gradient for small values of 𝒟 in Fig. 3**b**(ii).

### G. Propulsive hydrodynamic force generation

The leading order approximation to the hydrodynamic force generated by the swimming gait (i.e. the force required to hold the beating waveform fixed in place at the base), integrated over the arclength and averaged over a period of oscillation, can be calculated using resistive force theory (see Appendix B) to give

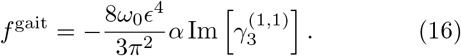

Note that this is a non-dimensional force (scaled by *ξ*^*n*^*L*^2^*/τ* ^*b*^), depending only on the beating kinematics and independent of the fluid viscosity [46]. This force is directed in the *x*-axis of the swimmer-fixed frame of reference. [17] showed that the swimming velocity due to a small oscillatory internal forcing scales with the beating frequency, the amplitude squared and the phase gradient. Our calculation for the hydrodynamic force *f* ^gait^ (proportional to the swimming velocity) due to self-organised chemomechanical oscillations agrees with this scaling relation—in this case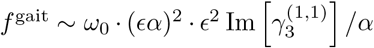. We can go further than previous results, however, by directly relating the hydrodynamic force to the chemomechanical parameters,

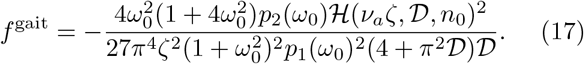

This expression is always negative, implying propulsion in the direction opposite to the base-to-tip propagating waves. Fig. 4 shows *f* ^gait^ as predicted by Eq. (17) and as calculated from simulations (again using RFT). The force increases with distance from the bifurcation, for all parameters (i)-(iv). Propulsion is enhanced for small diffusivity 𝒟 (panel (ii)), as expected since *f* ^gait^ inherits the 𝒟^−1^ dependence from the phase gradient. Lastly, *f* ^gait^ obtains a maximum as a function of the duty ratio *n*_0_, panel (iii), at a value slightly higher than the midpoint, mirroring the behaviour of the shear amplitude and phase gradient (Fig. 3**a**(iii) and **c**(iii)).

**FIG. 4.**
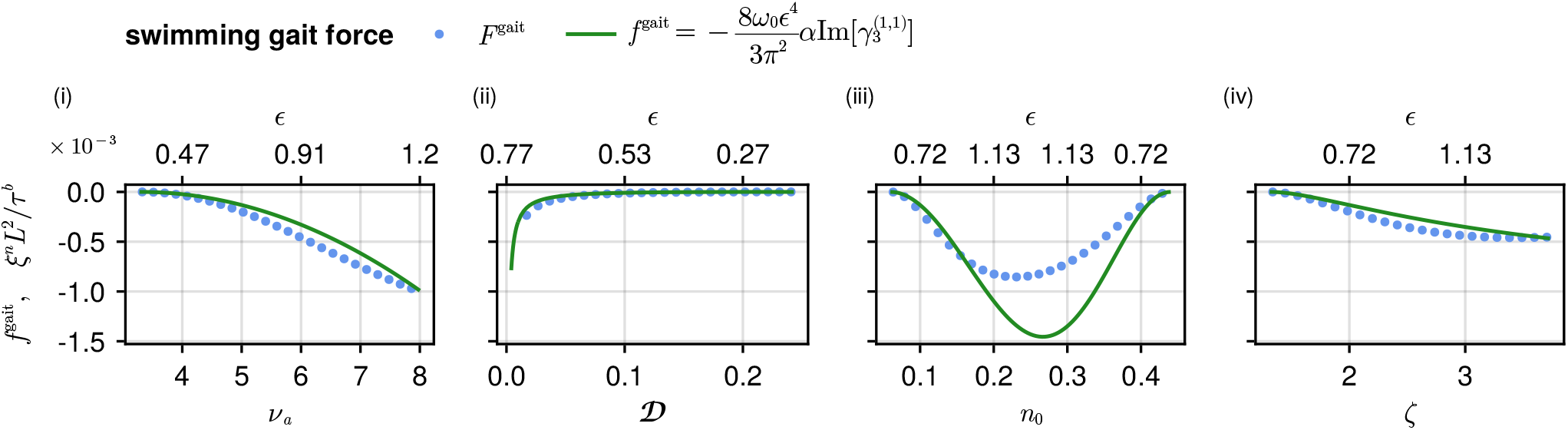
Weakly nonlinear prediction of hydrodynamic force generation of the swimming gait (green lines), compared with numerical simulations (blue markers). The time-averaged total force *f* ^gait^ (Eq. (17)), and numerically calculated force *F* ^gait^ (Appendix B), integrated over arclength *s*, in the head-fixed frame of the swimmer, using the same numerical simulations as in Fig. 3. Parameters are fixed at (*ν*_*a*_, 𝒟, *n*_0_, *ζ*) = (5.0, 0.02, 0.1, 2.0) if not explicitly varying.

### H. Motor binding distributions

The linear, 𝒪 (*ϵ*), solutions for the motor binding distributions read

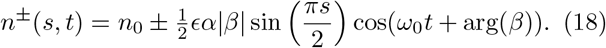

The two populations oscillate symmmetrically around the equilibrium value *n*_0_ in antiphase, with the same frequency *ω*_0_ as the shear angle *γ*. However the amplitude has an extra multiplicative factor where 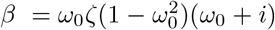 and the phase is offset by arg(*β*) = tan^−1^(1*/ω*_0_). Since 0 *< ω*_0_ *<* 1 the motors lead the shear angle by between a quarter and half of a cycle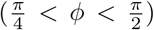. Fig. 5**a** shows the asymptotic solutions and numerical predictions for the saturation amplitude of differential motor activity *ñ*^sat^ (recall that *ñ* = *n*^−^ − *n*^+^). Eq. (18) can be understood, like Eq. (6), as the leading order term obtained from a spectral expansion of the differential motor activity,

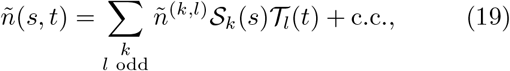

where *ñ*^(0,1)^ = *ϵαβ* + 𝒪 (*ϵ*^3^). We can then define *ñ*^sat^ := 2*ϵα*|*β*| as the saturing amplitude of differential motor activity. The tip amplitude max[*Ñ*(1)] (blue circles) and the numerical mode amplitude estimate 2|*Ñ*^(0,1)^| (orange circles) corresponding to the theoretical *ñ*^sat^ (solid green line), are shown in Fig. 5**a**. The predictions are again accurate for small to moderate values of *ϵ*, capturing the same qualititive variations as for *γ*^sat^ but with less quantitative accuracy for larger values of *ϵ*. Note that for the three parameters *ν*_*a*_, D and *ζ*, panels (i), (ii) and (iv), the tip amplitude max[*Ñ*(1)] approaches 0.2. This is expected because for an equilibrium value of *n*_0_ = 0.1, a difference of more than 0.2 would require the bound motor populations to become negative. This global con-straint (i.e. the motor populations must remain positive) is not captured at leading order in the expansion which assumes small amplitudes.

**FIG. 5.**
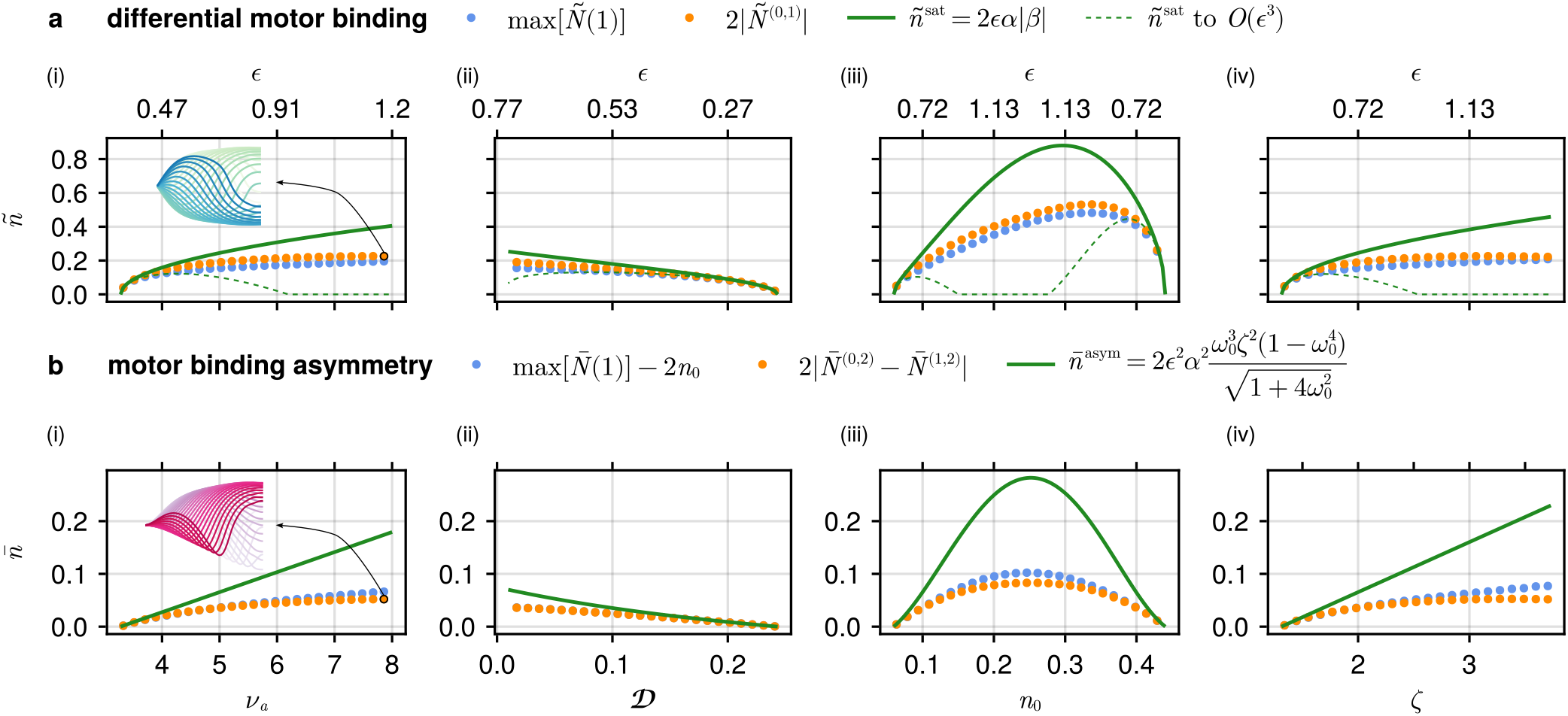
Weakly nonlinear predictions (green lines) of motor binding dynamics, compared with numerical quantities (blue/orange markers). Blue markers are quantities calculated directly from simulations, while orange markers are calculated from the spectral decomposition (Eq. (19)) of the simulation. Dashed green lines show predictions (generally only accurate for small values of *ϵ*) of one order higher in *ϵ* than solid lines. **a** Theoretical predictions of *ñ*^sat^ (Eq. (18)), the maximum difference in the fraction of attached motors on either side of the axoneme 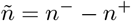 at *s* = 1, compared with the numerical maximum max[*Ñ*(1)] and spectral component 2|*Ñ*^(0,1)^. **a**(i, inset) The symmetric differential motor binding distribution *ñ*(*s, t*) over a cycle for the largest value of *ν*_*a*_. **b** Amplitude of the second harmonic 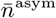 (Eq. (21)) of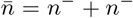, a measure of the asymmetry of motor binding oscillations about the equilbrium value, compared with the corresponding numerical quantities. **b**(i, inset) The asymmetric total motor binding distribution 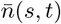 over a cycle for the largest value of *ν*_*a*_ simulated. Parameters are fixed at (*ν*_*a*_, 𝒟, *n*_0_, *ζ*) = (5.0, 0.02, 0.1, 2.0) if not explicitly varying.

The inset in panel **a**(i) shows the distribution of motor activity *ñ*(*s, t*) over a beating cycle as a function of arclength *s* for the largest value of *ν*_*a*_ simulated, highlighting the evolution from standing waves given by Eq. (18) at leading order in *ϵ* to propagating waves of motor activity away from equilibrium. These waves of motor activity, in turn, drive the propulsive beating patterns in the shear distribution.

### I. Asymmetry of molecular motor oscillations

Retaining terms at 𝒪 (*ϵ*^2^) in the perturbation expansion, a qualitative difference between the shear and motor oscillations emerges—asymmetric motor binding oscillations about the equilibrium value *n*_0_, due to the presence of the cos(2*ωt*) term oscillating at twice the fundamental frequency [47]. This is consistent with the system symmetry *γ* → −*γ, n*^*±*^ → *n*^∓^ (see Appendix A). Fig. 5**b** shows that the motor asymmetry increases with distance from the bifurcation. The inset beating pattern in panel **a**(i) visualizes asymmetric total motor distribution 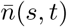 (as a function of *s*) over a cycle for the largest value of *ν*_*a*_ simulated. Since the maximum difference between motor populations *ñ* also increases with increasing distance from the bifurcation (Fig. 5**a**), the asymmetry of oscillations about the equilibrium value is a natural consequence of the fact that the motor populations cannot become negative. In other words, in order for the amplitude of motor binding oscillations to keep growing while remaining positive, they must become asymmetric. The total motor activity 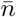 has an even expansion of the form

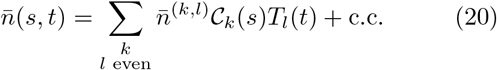

where *C*_*k*_(*s*) = cos(*kπs*) are the (even) shape modes. From this we predict an amplitude of the second frequency (see Appendix A for the detailed expansion coefficients) at *s* = 1 given by

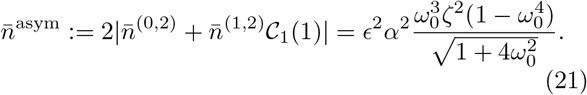

This prediction (green line) is compared against the maximum tip amplitude of the sum of motor populations, 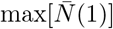, and the mode amplitude 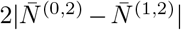 in Fig. 5**b**. As with the differential motor activity, the general trend is well predicted but the quantitative accuracy decreases with distance from the bifurcation. The expansions for the populations *n*^*±*^(*s, t*) are easily recovered from 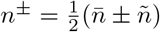 usingEq. (19) and Eq. (20).

### J. Spatial amplitude patterning

In our discussions of Fig. 3 to Fig. 5 we have compared asymptotic predictions and numerical solutions only at the tip of the flagellum (*s* = 1). However, the expansion Eq. (9) also predicts the spatial dependence of solutions. Fig. 6**a** shows modulations of the shear amplitude profiles [48]

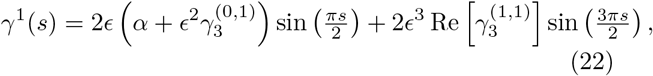

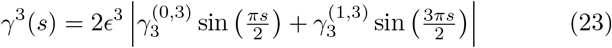

of the first and third temporal harmonics as *ϵ* increases away from equilbrium (see Appendix A for the explicit expansion coefficients). These expressions are calculated by the transformation taking Eq. (9) to Eq. (13). The functions of arclength *γ*^1^(*s*) and *γ*^3^(*s*) contain the shorter wavelength contribution S_1_(*s*) in their spectrum in addition to the fundamental standing wave component 𝒮_0_(*s*). The amplitude profile *γ*^1^(*s*) resembles a bullet shape with increasing *ϵ*; it becomes more flat towards the tip, with a relatively larger increase of amplitude nearer to the base. The third temporal harmonic *γ*^3^(*s*), in contrast, has a spatial profile with small amplitudes towards the base, and higher amplitudes towards the tip.

**FIG. 6.**
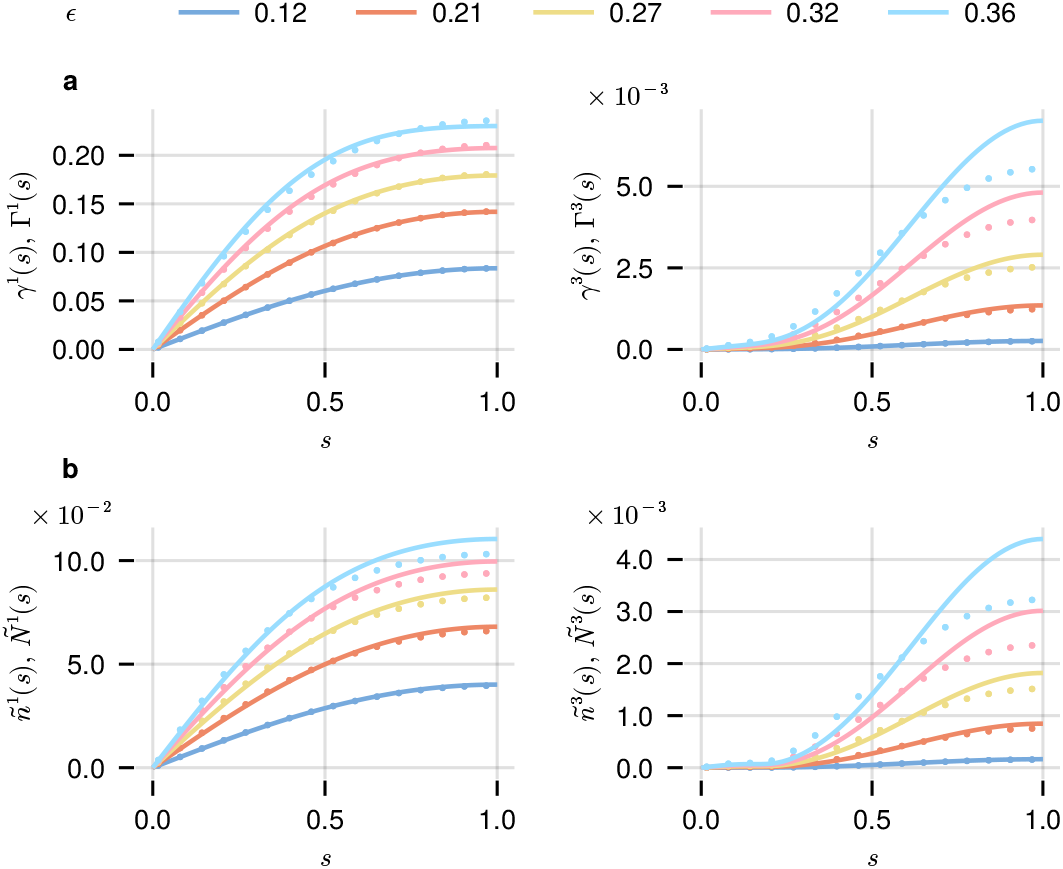
Modulation of numerical spatial amplitude profiles (markers) with increasing *ϵ* (by increasing *ν*_*a*_) compared with asymptotic predictions (solid lines) where third order 𝒪 (*ϵ*^3^) terms are retained. **a** Increasing amplitude of *γ*^1^(*s*) (left) and *γ*^3^(*s*) (right). The contribution of the third harmonic is more pronounced at the tip than close to the base. **b** Equivalent plots for the motor difference profiles *ñ*^1^(*s*) and *ñ*^3^(*s*).

For the parameter values used in Fig. 6**a** the third harmonic *γ*^3^(*s*), with leading order contribution 𝒪 (*ϵ*^3^), is seen to be roughly two orders of magnitude smaller than the first harmonic *γ*^1^(*s*). Less clear from the figure is how the 𝒪 (*ϵ*^3^) contribution to the third harmonic compares with the 𝒪 (*ϵ*^3^) contribution to the *first* harmonic, namely the coefficient *γ* of 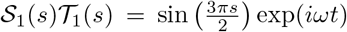 (hereafter we drop the dependence on *s* or *t* for standing wave modes). In fact this contribution is appreciable, which can be attributed to the 𝒟 ^−1^ dependence of the phase gradient and our selection in the numerics of 𝒟 = 0.02 (the third harmonic terms approach a constant as 𝒟 → 0). We discuss the 𝒟 dependence further in the following section. The same trends observed in Fig. 6**a** for the shear profile are also present in the differential motor population (Fig. 6**b**), where the functional forms of *ñ*^1^(*s*) and *ñ*^3^(*s*) are obtained in the same manner as Eq. (22) and Eq. (23), resulting in similar but slightly more complex expressions.

### K. Small diffusivity promotes the growth of shorter wavelength modes

7 explores the dependence of standing wave mode amplitudes on the diffusivity 𝒟. For *ϵ* = (i) 0.43 (ii) 1.0 and (iii) 2.39 we show the waveform predicted by

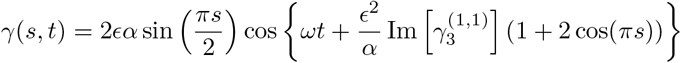

in column **a**, with 𝒟 = 0.02. From Tab. I, physical values are *D* ≈ 0.01 – 0.2, where smaller values are relevant for short ciliated organisms like *C. Reinhardtii*. The approximate solution in Eq. (24) agrees withEq. (6) to leading order, and both have an error of 𝒪 (*ϵ*^3^). Eq. (24), however, incorporates the correction term to the phase which comes from the phase difference between the oscillations of 𝒮_0_ and 𝒮_1_ at the fundamental frequency *ω* = *ω*_0_ +*ϵ*^2^*ω*_2_ (note that 𝒮_1_*/*𝒮_0_ = 1 + 2 cos(*πs*)). The corresponding numerical simulation is shown in **b** and the amplitude of the modes in the standing wave decomposition in **c**, again with 𝒟 = 0.02. Modes containing a first temporal mode, at frequency *ω* where *l* = 1, dominate the amplitude spectrum, consistent with Fig. 6. The two modes corresponding to a third temporal harmonic, 𝒮_0_𝒯_3_ and 𝒮_1_𝒯_3_, are much smaller than the 𝒮_1_𝒯_1_ term, although they all first appear at 𝒪 (*ϵ*^3^) in the expansion.

Interestingly, far from equilibrium the amplitude of 𝒮_1_𝒯_1_ can come to dominate the fundamental standing wave 𝒮_0_𝒯_1_ (**c**(iii)). In contrast, when 𝒟 is unity (only in column **d**) this effect is not present. The fundamental standing wave remains the most dominant, and all terms at 𝒪 (*ϵ*^3^) in the expansion are of comparable amplitude.

### L. Far from equilibrium beating patterns

So far we have obtained expressions from weakly nonlinear analysis that provide a biophysical understanding of the system dynamics close to the Hopf bifurcation point ℋ = 0, and even for *ϵ* ≈ 1. In this section we consider the far from equilibrium limit cycle dynamics for the parameters *ν*_*a*_ and *ζ*, for which the existence of oscillations is not bounded above (unlike 𝒟 and *n*_0_, see Fig. 3**a**). Hence we consider parameter values up to *ϵ* ≈ 4. Parameter values fit to experimental data suggest the biologically relevant regime consists of such far from equilibrium oscillations [12]. Although the asymptotic expansion is not theoretically valid in this regime, it is informative to examine the behaviour of numerical simulations and any deviations from predictions.

Fig. 8 shows the far from equilibrium behaviour of the system for variations in activity *ν*_*a*_ (left) and internal friction *ζ* (right). Superficially the frequency, phase gradient and hydrodynamic force behaviour (Fig. 8 **b**-**d**) appear very similar for variations in either parameter. These three plots show an initial linear regime that is predicted by the asymptotic expansion (green lines) with similar deviations in the numerical quantities (markers) thereafter. The shear amplitudes (Fig. 8**a**), however, trend oppositely, leading to qualitatively distinct waveforms (see **a**(i) and (ii) insets). Interestingly both the damped shear amplitude due to internal friction *ζ* and increasing shear amplitude with *ν*_*a*_ lead to the non-monotonic fall off of *f* ^gait^ (Fig. 8**d**), beyond the linear regime that is well predicted. From the asymptotics, Eqs. (7) and Eq. (8), we have that *γ*^sat^ decreases like 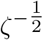 for large values of *ζ*. The non-monotonic dependence of the hydrodynamic force on *ν*_*a*_ and *ζ* is reminiscent of the dependence of propulsive force on the sperm number [46], when hydrodynamic moments are retained.

**FIG. 7.**
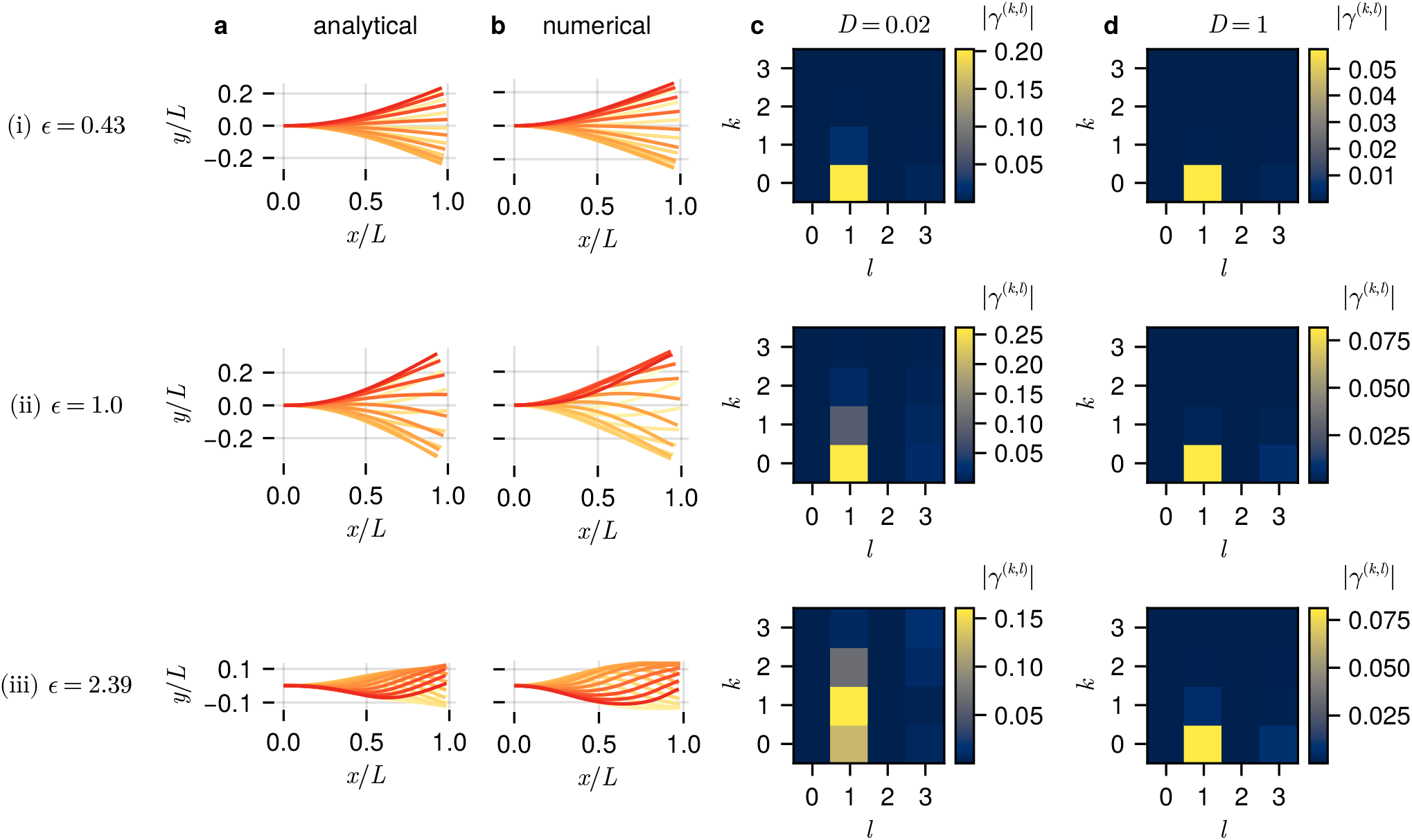
The waveform effect of the passive axoneme coefficient 𝒟 (diffusivity). Fig. 2-6 and **a**-**c** in this figure consider 𝒟 = 0.02, compared with **d** where 𝒟 = 1 i.e. the force required to bend the axoneme is comparable to the force required for shearing. **a** Asymptotic predictions (Eq. (24)) for three values (i)-(iii) of *ϵ* obtained by varying *ζ*. **b** Corresponding numerical simulations. **c** Spectral decomposition of the beat where the colorbar shows |*γ*^(*k,l*)^|, the amplitude of the mode 𝒮_*k*_(*s*) 𝒯_*l*_(*t*); (i)-(iii) show high contribution of the first harmonic (*l* = 1) compared with the third harmonic (*l* = 3). Panel **c**(iii) shows that the amplitude of the mode 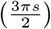 can become larger than 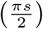 far from equilibrium. **d** Spectra for solutions where 𝒟 = 1, but *ζ* is tuned so that *ϵ* remains the same as in **a**-**c**. Here the 𝒮_0_(*s*)*T*_1_(*s*) mode always dominates, and the amplitude of the *S*_0_(*s*)*T*_3_(*s*) mode is comparable to the 𝒮_1_(*s*)*T*_1_(*s*) mode.

**FIG. 8.**
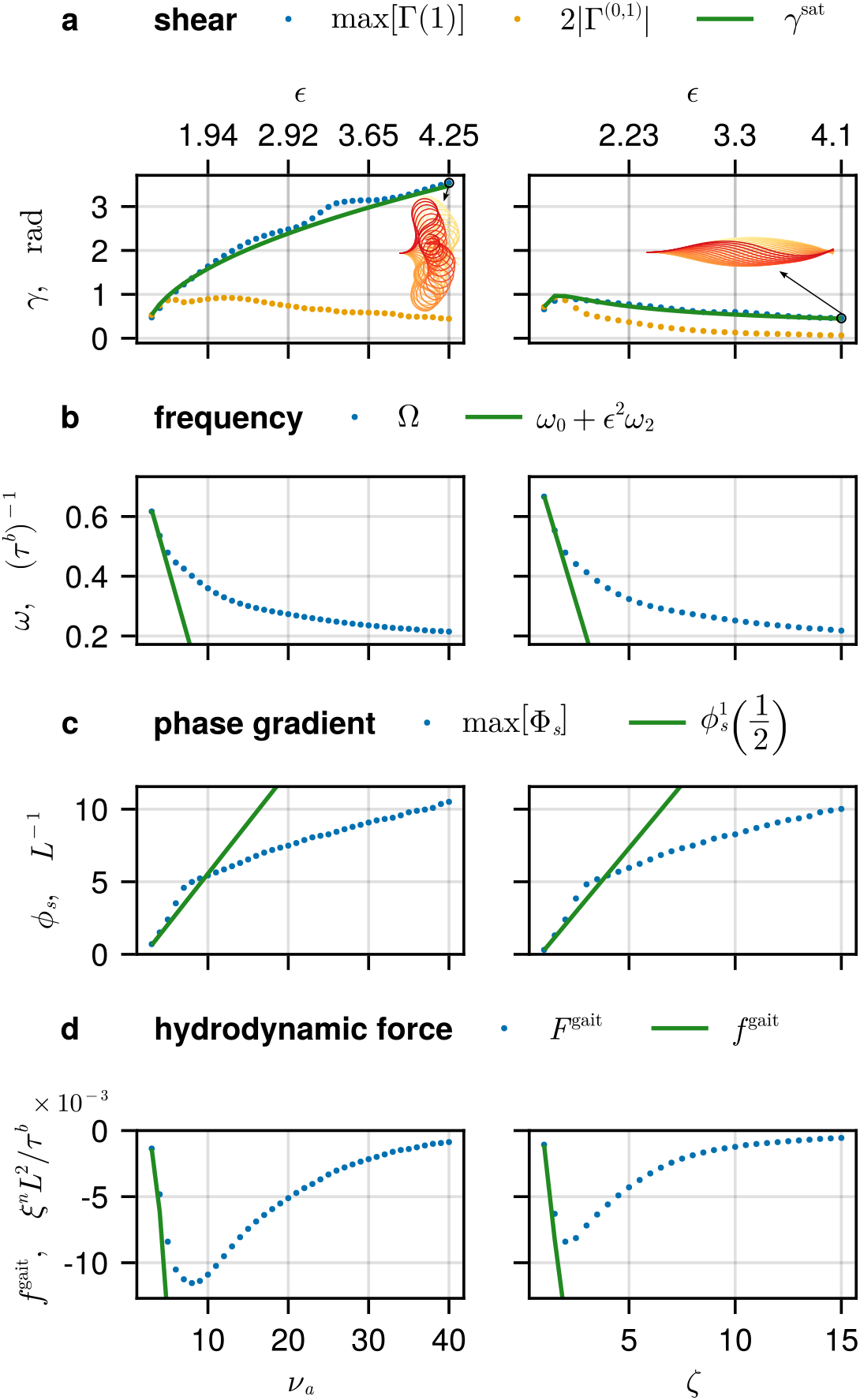
Far from equilibrium predictions (solid lines) vs. numerical quantities (markers), as functions of *ν*_*a*_ (left) and *ζ* (right). The same plots as Fig. 3 and Fig. 4 but for a larger range of *ϵ*, showing **a** the amplitude of shear *γ*, Γ, **b** frequency *ω*, Ω, **c** phase gradient *ϕ*_*s*_, Φ_*s*_ at the midpoint and **d** the gait force *f* ^gait^, *F* ^gait^. Although the effects of increasing *ν*_*a*_ and *ζ* on frequency, phase and propulsive force appear similar, the waveform modulations are quite distinct, see **a** inset beating patterns.

It is remarkable that the lowest order amplitude prediction, *γ*^sat^ = 2*ϵα*, i.e. the prediction of the amplitude of the fundamental standing wave mode (solid green line), remains close to the tip maximum max [Γ(1)] (blue circles) in this far from equilibrium regime, for both parameters. The calculated component of the fundamental standing wave Γ^(0,1)^ however, decays (orange circles). Roughly coincidental with the decay of the fundamental standing wave mode is an abrupt change of the slope in the variation of the phase gradient, and the turning point in the variation of the gait force.

## IV. DISCUSSION

Using a mutually validating combination of analytical and computational techniques we have uncovered new results related to sliding-controlled flagellar oscillations in terms of active and passive chemomechanical parameters of the flagellar-dynein system, for both weakly and strongly nonlinear beating patterns. We related these parameters, which describe nanoscale features of the flagellum interior, to microscale beat pattern observables via analytical expressions describing the amplitude (Eq. (7) and Eq. (8)), frequency (Eq. (12)), phase gradient (Eq. (14)) and hydrodynamic force generation (Eq. (17)). These expressions elucidate features of self-organised beating beyond the small-amplitude linear regime, leading to many novel insights that we summarise in the following.

### A. Oscillations are induced by a separation of time scales

The parameter combination *ν*_*a*_*ζ* = *ρf*_0_*/Kv*_0_*τ* ^*b*^ governs the transition to oscillations, with the onset controlled by a balance between active forcing, motor protein friction and passive bending and shear resistance. Hence, beyond the critical point, the timescale of viscoelastic shear relaxation *t*^*f*^ = *ρf*_0_*/Kv*_0_ is sufficiently longer than the timescale of relaxation of motor activity *t*^*b*^ = *π*_0_(*π*_0_ + *ϵ*_0_ exp(*f*^*^))^−1^. This is intuitive, for if the passive flagellum responds to perturbations quickly, the motors do not have time to generate the positive feedback required to drive the instability away from equilibrium. In terms of length scales, alternatively, the oscillations are controlled by the ratio of the displacement of an el-ement of the flagellum due to a team of motors against shear resistance *ρf*_0_*/K* to the average step length of a single unforced motor *v*_0_*τ* ^*b*^, highlighting the collective dynamics. Oscillations occur for only a limited range of values of the passive axoneme properties, quantified by diffusivity 𝒟 = *B/a*^2^*L*^2^*K*, and for the duty ratio *n*_0_ = *π*_0_*τ* ^*b*^ of motor attachment measured in the straight equilibrium configuration, while the existence of oscillations in the motor parameters *ν*_*a*_ = *ρf*_0_*/aK* and *ζ* = *a/v*_0_*τ* ^*b*^ is unbounded.

### B. Propulsive waves arise from standing wave mode interference

The asymptotic solutions reveal a decomposition of the beating pattern into a basis of standing wave modes, in which the solutions are low dimensional, consistent with other low dimensional flagellar representations in terms of principal components or asymmetry/parabolicity as discussed in previous studies [20, 35]. The principal shape components calculated for *C. Reinhardtii* beating recently [49, Fig. 1B] bear a strong resemblance to the modes 𝒮_*k*_(*s*) identified here. Here we have also used the standing wave basis to implement a spectral numerical method (see Appendix C), which is computationally inexpensive. Numerical simulations (see Fig. 7, for example) confirm the low dimensionality in this standing wave basis both near the equilibrium, where the weakly nonlinear analysis is valid, but also far from equilibrium where mode interactions are present.

The decomposition identified here may prove useful in the analysis of experimental waveforms, since it offers physical insights; most importantly, the nonlinear interaction of standing waves can produce a propulsive wave, required to generate net force over an oscillation period on the surrounding fluid. While the idea of superposed standing waves giving rise to travelling waves is familiar in the context of e.g. linear waves on a string, the nonlinear analogue in a dissipative system with interacting standing waves has not been explored in the cilia/flagella context previously. In our previous work we showed that nonlinear beating patterns of the RD model could be fit to experimental data for *C. Reinhardtii* cilia [12], in significant contrast to solutions of linearised models, for sliding-controlled oscillations [50]. We now have deeper insight into these beating patterns—away from equilibrium, where the solution is composed of many standing wave modes, non-trivial phase gradients can be generated. This effect is invisible at the linear level where the system goes unstable to a single standing wave mode 𝒮_0_𝒯_1_.

The phase difference generated depends in an important way on the (passive) diffusivity 𝒟 = *B/a*^2^*L*^2^*K*. Intuitively, as the bending elasticity gets larger, neighbouring elements are more tightly coupled and synchronous shear oscillations become more likely. As this coupling is weakened (in comparison with the local shear resistance), the flagellum is able to support higher frequency spatial modulations. The first spatial modulation at 𝒪 (*ϵ*^3^) comes from the mode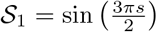, which, when oscillating out-of-phase with the fundamental mode 𝒮_0_, allows the emergence of propagating waves, with a phase difference that grows in magnitude with the distance from the equilibrium (Eq. (14)). We can express the diffusivity as 𝒟 = (*l*_*k*_*/L*)^2^, where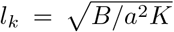. Intuitively, if the length of the flagellum is long compared to *l*_*k*_ i.e. 𝒟 is small, then the flagellum is able to support curvature that will be approximately on this scale. From the oscillator perspective, the more weakly coupled neighbouring sections are, the higher the wavenumber that can be generated. Physical estimates of 𝒟 (or *µ* = 𝒟^−1^) have tended to be small for *C. Reinhardtii* 𝒟 ∼ 0.1 and bull sperm D ∼ 0.01 [12, 41].

### C. Analytical expressions guide comparisons with experimental results

The variation of the amplitude with the flagellum length *L* is of critical interest, since *L* has been used as a control parameter in experiments; for example, [51] studied the variation of *C. Reinhardtii* waveform statistics as a function of *L*. A non-dimensionalisation that scales all forces with the bending force *B/L*^2^, as used in previous studies, obscures the effect of the length of the flagellum *L* on emergent waveforms. Here *L* appears only in the parameter 𝒟 = *B/a*^2^*L*^2^*K*, facilitating an easier comparison. We have seen in Fig. 3**a** that the shear tip amplitude is well predicted by 2*ϵα* (Eq. (7) and Eq. (8)). The only dependence on 𝒟 is in *ϵ*, which approaches a constant value 2*ν*_*a*_*ζn*_0_(1 − 2*n*_0_) − 1 as 𝒟 → 0 (or equivalently by increasing *L*). The amplitude should therefore reach a maximum value when 4 ≫ −*π*^2^D (see Eq. (7)) leading to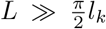, where again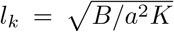. For previ-ously measured wild-type *C. Reinhardtii* parameters we estimate *l*_*k*_ ≈ 5.1 µm ([41], seeTab. I). Consistent with this prediction, Fig. 6**b** of [51] shows the maximum amplitude approaching a constant with increased length, at around the right scale (*L* ≈ 10 µm).

The frequency of oscillation depends only on the duty ratio *n*_0_ at leading order, with dependence on *ν*_*a*_*ζ* appearing with the 𝒪 (*ϵ*^2^) correction. Interestingly, there is no dependence of frequency on the diameter *a* of the flagellum which drops out in the combination *ν*_*a*_*ζ*. We have seen that the frequency tends to decrease as the system moves away from equilibrium (see Fig. 3**b**). The decreasing frequency with motor activity *ν*_*a*_ has been discussed previously [20, 26], where it was suggested that an increased density of motors with the same attachment rate requires more motors to unbind to affect changes in the flagellar shape, leading to delays compared with microtubules that are more sparsely populated with motors. Experimentally, the concentration of ATP/ADP in solution around a demembranated axoneme is often varied [34, 35]. Since ATP is required for detachment of dynein from microtubules, we may expect a decrease in the equilibrium duty ratio *n*_0_ = *π*_0_*/*(*π*_0_ + *ϵ*_0_ exp(*f*^*^)^−1^), which scales inversely to the base detachment rate *ϵ*_0_, when the concentration of ATP is increased. Fig. 3**b**(iii) shows the model predicts increasing frequency with decreasing *n*_0_ (increasing ATP), consistent with experimental findings [34, 35]. Increasing the length *L* (corresponding to decreasing 𝒟), on the other hand, leads to a slowly decreasing frequency. This is potentially consistent with the behaviour observed in [51, Fig. 6**a**], although there the frequency appears to increase slightly before decreasing.

We calculated the hydrodynamic force generated by the beating pattern in the body fixed frame of reference (Eq. (17)), equivalent to the force required to clamp the beating flagellum to a wall at the base in the lab frame. We find once more that the predicted increase in force with length (Fig. 4) matches the experimental result [51, Fig. S3**A**]. This force is directed oppositely to the base-to-tip propagating waves. We showed that the force scales with the waveform parameters as ∼ *α*^2^*ω*∂_*s*_*ϕ*, consistent with earlier work [17]. This leading order approximation captures the increase in force near the bifurcation (Fig. 4). Fig. 8 shows non-monotonic behaviour, however, further from equilibrium (this may be captured by the next order correction to the force, which is 𝒪 (*ϵ*^6^), and is not included in the asymptotic analysis here).

The fact that the total hydrodynamic force scales with ∼ *ϵ*^4^ provides a consistency check on neglecting the contribution of hydrodynamic forces (in low viscosity) to the internal moment balance (Eq. (3)) when compared with the 𝒪 (*ϵ*) shear resistance term *γ*. The leading order hydrodynamic forces (using slender body theory or RFT [52]) scale with ∼ *ξ*^*n*^*L*^2^*ω*, where the resistive force coefficient in the direction perpendicular to the flagellum *ξ*^*n*^ is proportional to the fluid viscosity. High viscosity, therefore, may generate forces that have a nonnegligible effect on the beating pattern, taking us out of the reaction-diffusion regime considered in this work. For instance, hydrodynamic moments acting non-uniformly along the flagellum can induce higher wavenumbers and buckling instabilities [53, 54].

Predictions of observables as functions of parameters other than *L*, as we have done in this section, could also be tested in this manner. For example, we predict optimal values of the duty ratio *n*_0_ for the shear amplitude (Fig. 3(iii)) and hydrodynamic force (Fig. 4(iii)). At present, however, the motor binding times (and their response to load) of axonemal dynein are not measurable in a beating cell. We hope that this work and further theoretical elaborations may inform experimental work in this area.

### D. The fundamental standing wave may power propulsive waveforms in low viscosity

The amplitude expression, Eq. (8), for the fundamental standing wave mode 𝒮_0_(*s*) 𝒯_1_(*t*) predicts the saturating amplitude *γ*^sat^ and mode amplitude 2|Γ^(0,1)^| accurately in the weakly nonlinear regime, and remarkably continues to predict the tip amplitude max[Γ(1)] even for highly nonlinear solutions. We saw, however, in Fig. 8**a** that while the theoretical prediction of the amplitude remains accurate, the actual amplitude |Γ^(0,1)^| (orange circles, calculated from simulations) of the fundamental standing wave decays, and this decay correlates with decreasing hydrodynamic force (Fig. 8**d**). This suggests a strong fundamental standing wave may be a necessary component of a propulsive waveform; while not propulsive in itself, it provides the background against which other modes generate asymmetry and propulsion. A standing wave component to the beat of *C. Reinhardtii* was also indicated by the analysis of [49].

For large values of *ν*_*a*_, the decay of the fundamental standing wave suggests the energy provided by the motors must go into transverse (reciprocal) oscillations rather than tangential (non-reciprocal) oscillations that would generate a directed gait force. It is not obvious a priori what the effect of large *ζ* will be on the system dynamics. On one hand, increasing *ζ* increases internal viscous drag, resulting in smaller sliding velocities. On the other hand, increasing *ζ* changes the force that motors experience, thus increasing the responsiveness to changes in sliding velocity, which appears in the exponential term in Eq. (4). The non-monotonic dependence of the amplitude on *ζ* may therefore reflect the importance of the detachment response at low values and of increased friction at high values (Fig. 8**d**).

### E. Limitations

The benefit of considering a simple reaction-diffusion model to describe flagellar beating is that, as we have seen, we can make theoretical predictions that go beyond linearisations and thus bring the model into closer contact with the out-of-equilibrium nature of experiments. Of course, by simplifying we also restrict when the model might usefully be applied. We cannot make predictions regarding changes in viscosity using this model, for instance, which would be available as an experimental control parameter. In fact, simulations of the corresponding chemo-elastohydrodynamic system (the RD model extended to include hydrodynamic forces using RFT) have been unable to capture high viscosity experimental waveforms such as those displayed by spermatozoa [12, 55], suggesting modifications may be necessary.

Measurements of the duty ratio *n*_0_ at the stall force of axonemal dynein would be valuable data that could be used to predict experimental observables, but it is not clear that single molecule measurements of the pro-portion of time spent attached by the motor under load would be equivalent to the model quantity *n*_0_[56], since the model coarse-grains the dynein cycle into two states—attached (force producing) and detached (not force producing). Dynein is well known to cycle through more than two conformations (see [57] for review) and there is some debate around which of these are force producing [58, 59].

We have not considered intrinsically asymmetric waveforms, as is the case for wild-type *C. Reinhardtii*. It is straightforward to add an intrinsic curvature to the axoneme around which the model may describe oscillations [25], but it remains a challenge to reproduce the behaviour observed in [60] in a model system, where increasing activity initially increases static curvature, before a dynamic instability superposes oscillations.

### F. Outlook

Ciliated organisms are multi-scale systems [11]. At the subcellular level, small populations of motors engaged in a tug of war can generate local flagellar oscillations. As we have seen, weak coupling of oscillating units through bending of flexible microtubules enables a phase-gradient or wavenumber to emerge that enables propulsive swimming or fluid pumping. From here, there are many possibilities for extensions; for example, these ideas are repeated at the level of an organism with many cilia such as paramecium, or for populations of cilia lining the human respiratory tract. Dynamics of many cilia systems can be described by a population of robust oscillators that are weakly coupled, at this scale via hydrodynamic or elastic interactions through a substrate, that self-organise into metachronal waves [10, 31, 61, 62]. The reaction-diffusion system then becomes a single oscillatory unit at this larger scale, and the same techniques may be applied. Other extensions to the fundamental framework could include changing the motor binding kinetics or implementing three-dimensional beating. We provide the code for the symbolic mathematics used to generate the expansion and the spectral numerical method at github.com/polymaths-lab. Similar analyses applied to other models of self-organised flagellar oscillations, such as the geometric clutch [16, 18], could provide predictions that may be used to distinguish between models.

The unknown mechanism of flagellar locomotion remains a key medical question, since ciliary dysfunction is related to fertility issues and a number of diseases [63]. While it is still unclear that the sliding-controlled mechanism [12, 17, 20, 28, 31, 36] is the one used by cells, the results presented here provide a framework for further investigation, and a number of testable predictions. The principles described here may also be applied to the design of biomimetic systems.

## Appendix A Weakly nonlinear analysis

We here outline the asymptotic method by which the results were obtained (the full derivation using the computer algebra package Sympy is also available in a Jupyter notebook at github.com/polymaths-lab).

1. First we shift *n*^*±*^ such that the equilibrium value (*γ, n*^*±*^) = (0, *n*_0_) is at the origin, and make the change of variables *ñ* = *n*^−^ − *n*^+^ and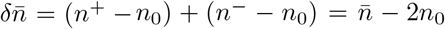, where 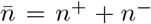. Eq. (3) and Eq. (4) are recast with respect to these new variables:

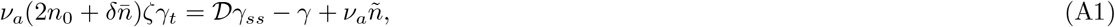

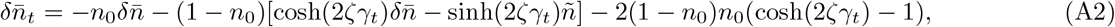

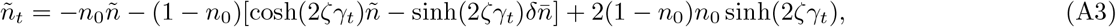

where we have set *f*^*^ = 2.

2. We introduce a small parameter 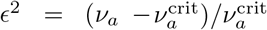 which measures the relative distance from the point of Hopf bifurcation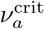. Any solution must remain a solution under the transformation *ϵ* → −*ϵ* (having no effect on the governing equations), which can be enforced using the system symmetry which interchanges the motor populations *n*^+^ ↔ *n*^−^ and sends *γ* → −*γ*. This leads to perturbation ansatzes *γ* = *ϵγ*^(1)^ + *ϵ*^3^*γ*^(3)^ + *O*(*ϵ*^5^), *ñ* = *ϵñ*^(1)^ + *ϵ*^3^*ñ*^(3)^ + *O*(*ϵ*^5^) which include only odd terms in *ϵ* and hence change sign under the symmetry, and 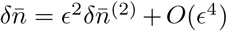 with only even terms that remain invariant.

3. We use Lindstedt’s method [64] in order to allow the frequency *ω* = *ω*_0_ + *ϵ*^2^*ω*_2_ + *O*(*ϵ*^4^) to depend on *ϵ*. This involves scaling time by *ω*^−1^ before substituting the expansions.

4. We match terms at each power *ϵ*^*j*^, obtaining a system of two linear equations for the unknowns **u**^(**j**)^ = [*γ*^(*j*)^ *ñ*^(*j*)^ ]^**T**^ at odd orders of the form

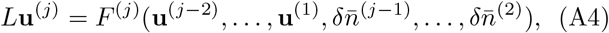

and a single equation for 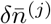 at even orders. The matrix-differential linear operator *L* reads

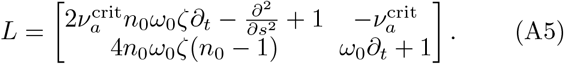

5. The linear solutions of *L***u**^(**1**)^ = **0**, obained at *O*(*ϵ*), are given by

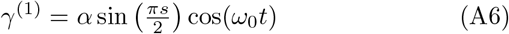

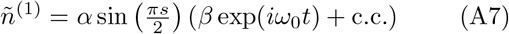

where the amplitude *α* is arbitrary, and we have chosen the arbitrary phase (due to the continuum of limit cycle solutions) of the cosine to be zero. The critical frequency is found to be 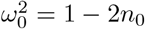 and *β* = 2*n*_0_*ω*_0_*ζ*(*ω*_0_ + *i*) gives the amplitude and relative phase of the motor oscillations to the shear. The critical activity value reads

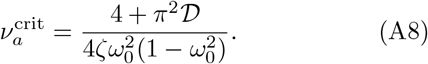

6. The solution of the single linear equation at *O*(*ϵ*^2^) is given by

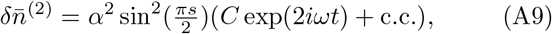

where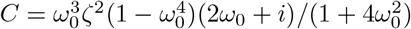.

7. Substituting the solutions obtained so far into the *O*(*ϵ*^3^) equations gives

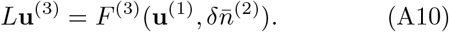

Now we apply the Fredholm alternative theorem [65]: for solutions to exist at *O*(*ϵ*^3^), the right hand side must be orthogonal, with respect to the complex inner product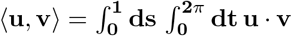, to the null space of the adjoint operator *L*^*†*^ of the linear operator *L*. Applying this solvability condition we determine

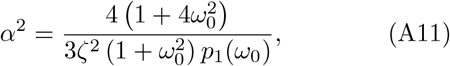

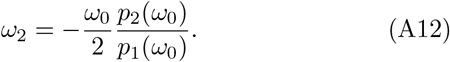

where 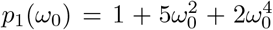 and 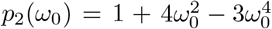.

8. With these choices of *α* and *ω*_2_ substituted into the right hand side function *F* ^(3)^, it is now possible to solve the *O*(*ϵ*^3^) equations, which split into linear systems whose coefficients are products of the spatial and temporal Fourier modes 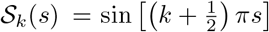 and 𝒯_*l*_(*t*) = exp(*liωt*) for *k, l* = 0, 1, 2, These are solved using the method of undetermined coefficients. In the same manner as in step 5, there is an arbitrary multiplicative constant *α*_3_ of the fundamental standing mode 𝒮_1_(*s*) 𝒯_1_(*t*) coming from the homogeneous problem, which appears at 𝒪 (*ϵ*^3^). Full solutions follow in Appendix A. Repeat steps 6-8 to obtain the solutions at *O*(*ϵ*^4^) and *O*(*ϵ*^5^) and to fix the arbitrary amplitude *α*_3_.

## Appendix B Complex Expansion Coefficients

The expansion of the shear angle *γ*(*s, t*) up to an error of 𝒪 (*ϵ*^5^) is given by

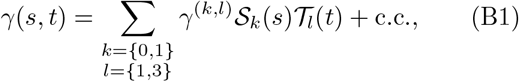

where

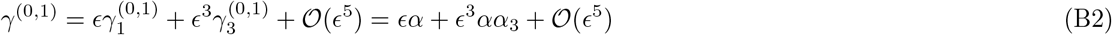

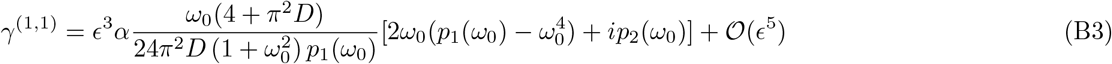

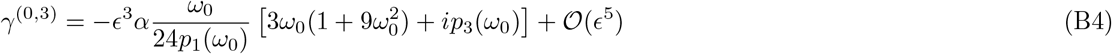

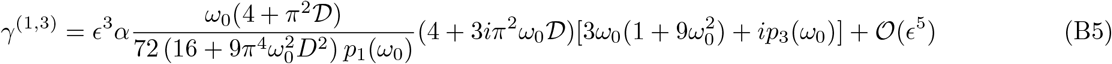

where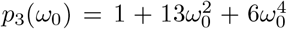. The amplitude *α*_3_ that appears at 𝒪 (*ϵ*^3^) is arbitrary at this level. By going to the next order we find *α*_3_ to consist of a twelth order polynomial in *ω*_0_ with coefficients that are cubics in 𝒟. This expression is not given here, but can be found in the accompanying Jupyter notebook github.com/polymaths-lab. The expansion of the differential motor population *ñ*(*s, t*) = *n*^−^(*s, t*) − *n*^+^(*s, t*) is given by

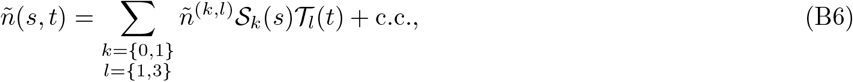

where

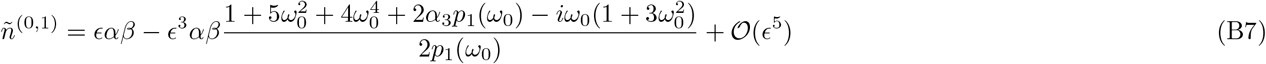

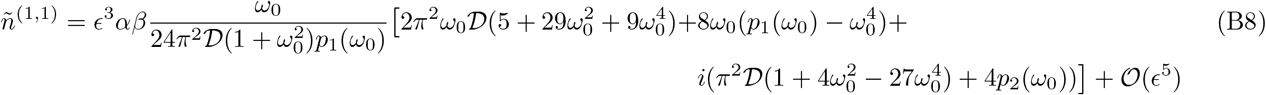

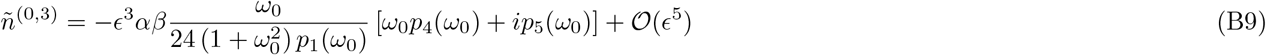

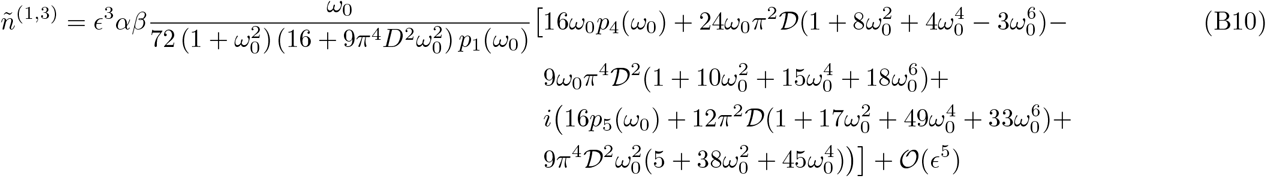

where 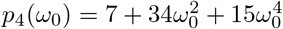 and 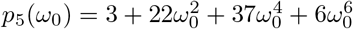.

Finally, the expansion of the total motor population 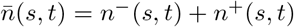 is given by

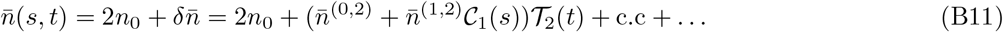

since 𝒞_0_(*s*) = 1, where

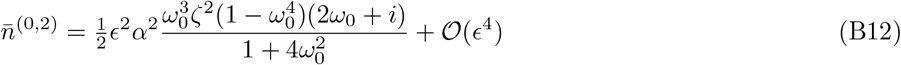

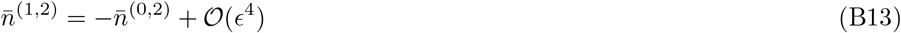

## Appendix C RFT Calculation of the Propulsive Force

The total hydrodynamic force on the flagellum is given by resistive-force-theory (RFT, [52]) as

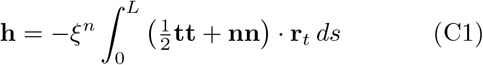

where *ξ*^*n*^ is the drag force per unit length against motion normal to the flagellar centerline, and tangential drag is taken to be half the size (a common approximation). In non-dimensional form (with hydrodynamic force scaled by *ξ*^*n*^*L*^2^*/τ*) this reads

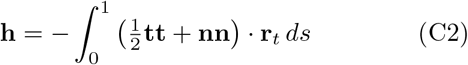

Since **r** = ∫^*s*^ (cos(*γ*), sin(*γ*)) *ds*^*′*^, we may plug in the asymptotic expression for *γ* including the leading order phase correction, Eq. (24), and expand all transcendental functions. Finally integrating over a period *T* we obtain the leading order gait force, Eq. (17). Numerically we useEq. (C2) with the simulated Γ, integrating over the arclength with the trapezoidal rule.

## Appendix D Numerical Methods

To obtain accurate limit cycle solutions numerically, we employ a predictor-corrector method. The predictor solves the initial value problem given by Eq. (3) and Eq. (4) with a discretized (N=63 points) initial shear distribution given by a small Gaussian perturbation from the straight equilibrium state *γ* = 0 (with the motor populations at equilibrium *n*^*±*^ = *n*_0_), as detailed in our previous work [12].

After converging to a solution close to the limit cycle, the corrector uses one cycle of the predictor solution and the approximate period *T* as an initial guess for a Newton-Raphson method, which is used to solve the boundary value problem given by Eq. (3) and Eq. (4) augmented with periodic boundary conditions:

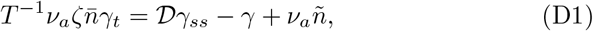

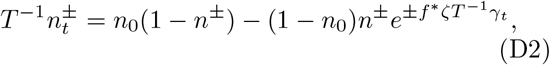

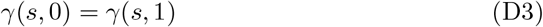

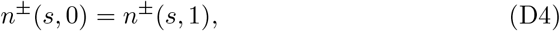

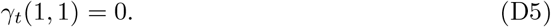

In this method, time is first scaled by the unknown period *T*, which becomes an extra unknown in the rootfinding problem. Derivatives are performed in frequency space, whereby the discrete Fourier transform turns temporal derivatives into multiplication, and automatically enforces periodicity in time. Spatial derivatives *γ*_*ss*_ are conveniently implemented by a discrete sine transform (DST) of type S3, which decomposes a signal into exactly the modes 𝒮_*k*_(*s*) used in the asymptotic expansion (see Appendix A). The second derivative is then obtained in frequency space by a multiplication with (minus) the square of the spatial wavenumber *k*. The inverse of this transform is also a DST, of type S2. The corrector is iterated until the residual is below a relative tolerance of 10^−6^. Finally, the phase condition *γ*_*T*_ (1, 1) = 0 picks out the particular periodic solution where the tip of the flagellum is at a maximum/minimum at the end of the cycle, providing an extra equation corresponding to the extra unknown *T*.

The above scheme is implemented in the Julia language using the DifferentialEquations.jl package [66]. The code is available at github.com/polymaths-lab.

